# In vivo human whole-brain Connectom diffusion MRI dataset at 760 μm isotropic resolution

**DOI:** 10.1101/2020.10.05.327395

**Authors:** Fuyixue Wang, Zijing Dong, Qiyuan Tian, Congyu Liao, Qiuyun Fan, W. Scott Hoge, Boris Keil, Jonathan R. Polimeni, Lawrence L. Wald, Susie Y. Huang, Kawin Setsompop

## Abstract

We present a whole-brain in vivo diffusion MRI (dMRI) dataset acquired at 760 μm isotropic resolution and sampled at 1260 q-space points across 9 two-hour sessions on a single healthy subject. The creation of this benchmark dataset is possible through the synergistic use of advanced acquisition hardware and software including the high-gradient-strength Connectom scanner, a custom-built 64-channel phased-array coil, a personalized motion-robust head stabilizer, a recently developed SNR-efficient dMRI acquisition method, and parallel imaging reconstruction with advanced ghost reduction algorithm. With its unprecedented resolution, SNR and image quality, we envision that this dataset will have a broad range of investigational, educational, and clinical applications that will advance the understanding of human brain structures and connectivity. This comprehensive dataset can also be used as a test bed for new modeling, sub-sampling strategies, denoising and processing algorithms, potentially providing a common testing platform for further development of in vivo high resolution dMRI techniques. Whole brain anatomical T_1_-weighted and T_2_-weighted images at submillimeter scale along with field maps are also made available.

## Background & Summary

Diffusion magnetic resonance imaging (dMRI)^1^ has proven to be an invaluable tool in the study of brain structural connectivity and complex tissue microarchitecture. It has also offered the ability to investigate a variety of neurological, neurodegenerative and psychiatric disorders^2–10^. In recent years, a number of dMRI data repositories^11–18^ have been published to enhance the understanding of human brain, including large-scale population study projects launched across the world, such as the Human Connectome Project (HCP)^19–22^ and the UK Biobank project^23^. However, most of these studies acquired in vivo dMRI datasets at a relatively low spatial resolution of 1.25 - 3 mm, due to technical difficulties in achieving higher spatial resolution that include: i) the inherent low SNR of dMRI acquired with smaller voxels; ii) the distortion/blurring artifacts from longer EPI readouts; and iii) the inevitable subject motion during long scan time required for obtaining multiple diffusion directions. Submillimeter isotropic resolution dMRI datasets have therefore been mostly confined to ex-vivo studies^24–26^ where there are no scan time and motion constraints. While ex vivo dMRI plays an important role in studying brain structures with its higher achievable resolution, the alteration of tissue properties due to fixation may fundamentally change the dMRI measurement and affect its ability to reflect accurate values in the living human brain. In vivo dMRI serves as a non-invasive probe of tissue structure in different brain pathologies, and can also be combined with functional and behavioral data for further investigation of structure-function relationships. The availability of submillimeter resolution in vivo dMRI datasets would enable more detailed examination of the brain’s fine-scale structures, such as probing short-association fibers at the gray-white matter boundary^27^ or studying the cytoarchitecture of thin cortical layers (< 1-4 mm in thickness)^26,28^. Submillimeter resolution in vivo dMRI would also facilitate the development and optimization of new sampling, modeling and processing methods specifically targeted at recovering information about tissue microstructure from such high resolution in vivo dMRI datasets.

With the advent of advanced hardware^29–31^, acquisition and reconstruction strategies^32–37^, submillimeter isotropic resolution dMRI is becoming increasingly feasible^38–42^, presenting new opportunities for studying brain structure at the mesoscopic scale in vivo. In this work, we aim to create the first publicly available in vivo whole brain dMRI reference dataset acquired at submillimeter isotropic resolution of 760 μm, i.e. at 7.7 times smaller voxel size than conventional dMRI acquisition at ~1.5 mm resolution. To enable high-SNR at this extreme resolution, state-of-the-art hardware and acquisition techniques were used along with a carefully designed diffusion protocol targeting at a single-subject scan across 9 two-hour sessions. The data were acquired on the MGH-USC 3T Connectom scanner equipped with high-strength gradient (G_max_ = 300 mT/mp)^29,30,43^ and a custom-built 64-channel phased-array coil^31^, using the recently developed SNR-efficient simultaneous multi-slab imaging technique termed gSlider-SMS^40,41^. To mitigate potential image artifacts and loss in resolution, a custom-made form-based headcase (https://caseforge.co) that precisely fit the shape of subject’s head and the coil was also used to ensure minimal motion contamination, consistency of subject positioning and same field shim settings across sessions. Advanced parallel imaging reconstruction with ghost-reduction algorithm (Dual-Polarity GRAPPA^44,45^) and reversed phase-encoding acquisition were also performed to mitigate ghosting artifacts and image distortion. We acquired a total of 1260 q-space points including 420 directions at b=1000 s/mm^2^ and 840 directions at b=2500 s/mm^2^ across the 9 sessions to ensure high angular resolution. An optimized preprocessing pipeline was employed afterwards to correct for intensity drifts, distortions and motion while preserving as much high-resolution information as possible.

We envision that this reference dMRI dataset will have a broad range of investigational, educational, and clinical applications that will advance the understanding of human brain structure. In particular, the data should be useful in the exploration of mesoscale structures revealed through submillimeter in vivo dMRI, and in the evaluation of structural brain connectivity that should improve with reduced intravoxel fiber complexity using smaller voxels and with higher intravoxel fiber crossing resolution using high angular resolution^46^. The data may also serve as a test bed for new modeling, data subsampling, denoising and processing algorithms, potentially providing a common testing platform for further technical development of in vivo high resolution dMRI.

## Methods

### Participant

The participant was a young healthy Caucasian man (born in 1989). He was fully instructed about the scans and gave written informed consent for participation in the study. The experiments were performed with approval from the institutional review board of Partners Healthcare.

### Data acquisition

All data were acquired in 9 two-hour scan sessions on the MGH-USC 3T Connectom scanner equipped with a maximum gradient strength of 300 mT/m and a maximum slew rate of 200 T/m/s ^29,30,43^, using a custom-built 64-channel phased-array coil ^31^. A custom-designed personalized motion-robust stabilizer (https://caseforge.co), that was precisely fit to the shape of the subject’s head and the inside of the coil, was used to ensure the same subject positioning across sessions and minimal motion contamination within sessions (Fig. 1a). Field shim settings were kept identical across sessions to minimize differences in image distortion, with the goal of minimizing reliance on image post-processing that could introduce blurring.

**Figure 1.**
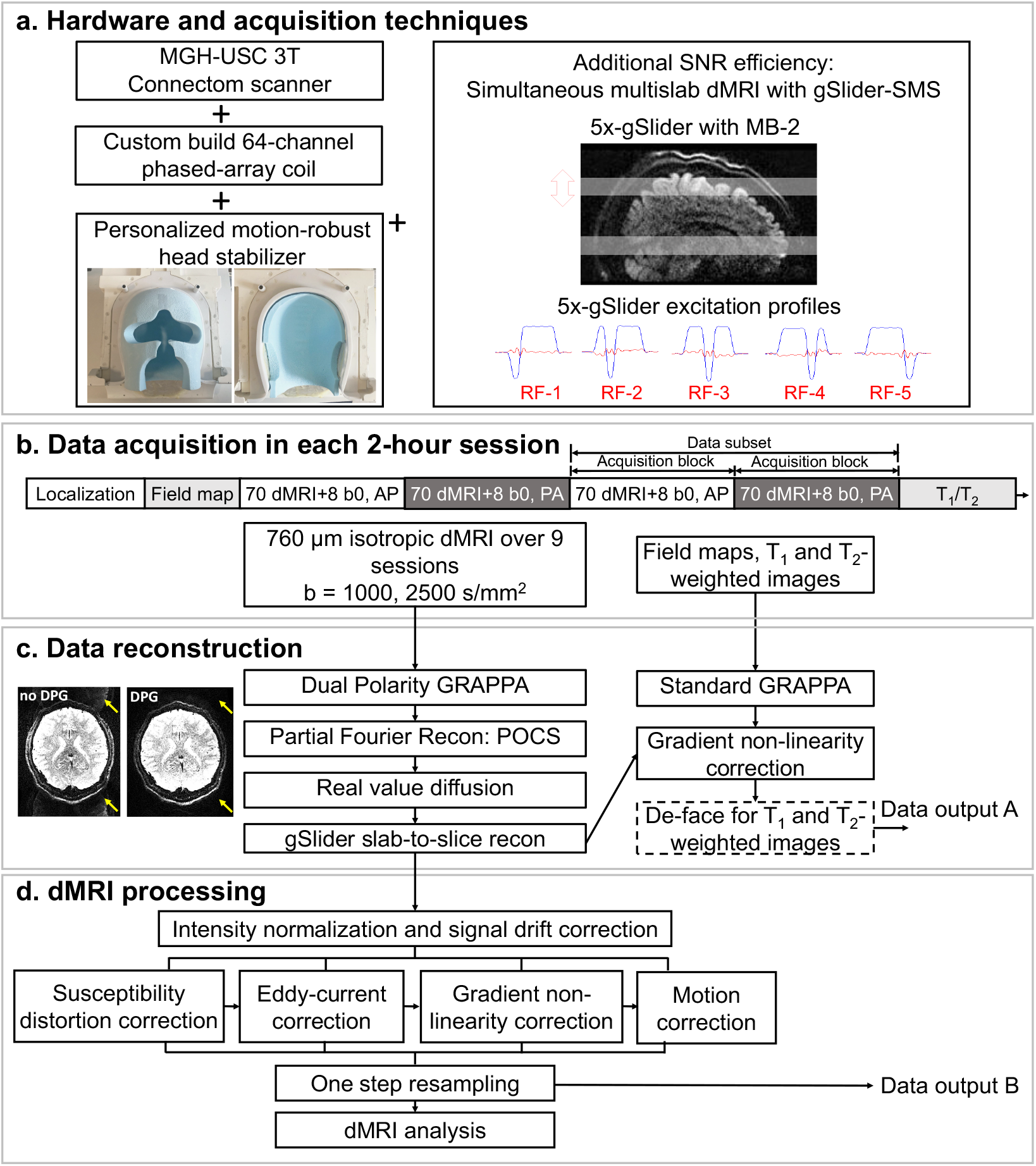
Data acquisition, reconstruction and preprocessing workflow. (a) Hardware and acquisition techniques used to acquire the data. The Connectom scanner, the custom-built 64-channel phased-array coil and the gSlider-SMS acquisition technique provide high SNR, while the personalized motion-robust stabilizer ensures consistent positioning across multiple scan sessions and minimal motion contamination within each session. (b) Data acquisition pipeline in each 2-hour session. The dMRI acquisition was divided into 4 acquisition blocks, each of which comprised 70 diffusion volumes and 8 interleaved b = 0 s/mm^2^ volumes. Two acquisition blocks with paired reversed phase encoding composed a data subset with a unique set of diffusion directions. 6 data subsets were acquired at b = 1000 s/mm^2^ in the first 3 sessions, and 12 data subsets at b = 2500 s/mm^2^ in the remaining 6 sessions. (c) Data reconstruction workflow. dMRI data were reconstructed using Dual Polarity GRAPPA, partial Fourier reconstruction, real value diffusion and gSlider slab-to-slice reconstruction, while field maps and structural images were reconstructed using standard GRAPPA reconstruction implemented on the scanner. Minimally preprocessed data were then provided after the gradient non-linearity correction (Data output A). (d) dMRI preprocessing with one-step resampling. The preprocessed data were obtained after intensity normalization to correct for signal drift, along with susceptibility-induced and eddy current-induced distortion correction, gradient non-linearity correction and motion correction, all of which were performed together using a one step resampling process to minimize interpolation-induced image blurring (Data output B).

To obtain high spatial resolution dMRI data, the gSlider-SMS^40^ sequence (Fig. 1a) was used in our study. gSlider-SMS is a recently developed self-navigated simultaneous multi-slab acquisition that combines blipped-CAIPI^47^ controlled aliasing with a slab-selective radio frequency encoding, which has been demonstrated to provide high SNR-efficiency for high resolution dMRI. The gSlider-SMS protocol to acquire these data was constructed to achieve an isotropic submillimeter resolution with whole brain coverage, high SNR, minimal reconstruction errors, reduced distortion and blurring as well as tolerable scan time per volume. Motivated by this goal, the decision of the final imaging parameters was made by evaluation of data quality of pilot scans and conclusions from prior studies. The 0.76-mm isotropic resolution was chosen as it provides a good trade-off in term of resolution and scan time to achieve reasonable SNR. A gSlider encoding factor of 5 and a multi-band acceleration factor of 2 were used to achieve high SNR efficiency by acquiring 10 imaging slices simultaneously per EPI-readout across two thin RF-encoded slabs. In-plane acceleration of 3 was selected to mitigate EPI distortion and blurring by reducing the effective echo spacing from 1.02 ms to 0.34 ms, while ensuring good reconstruction performance with low noise amplification when the reconstruction was performed with Dual polarity GRAPPA to also mitigate ghosting artifacts. The detailed imaging parameters are listed in Figure 2a.

**Figure 2.**
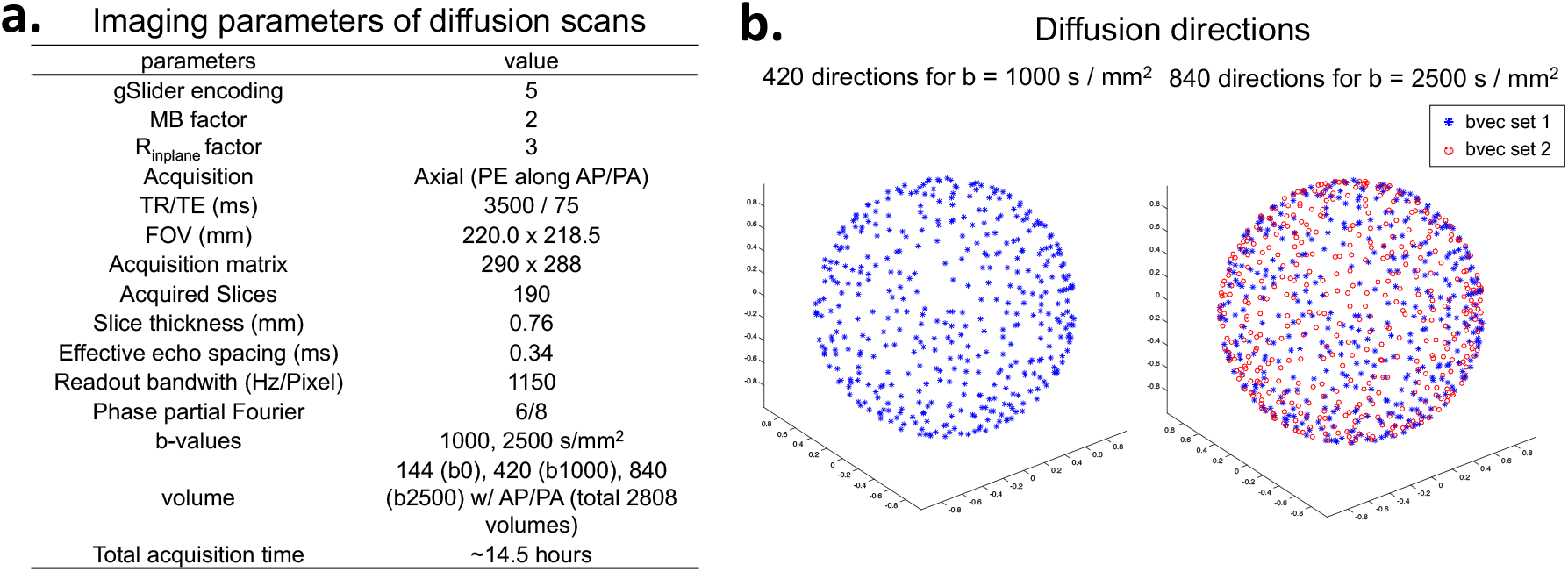
Imaging parameters for the dMRI dataset. (a) Detailed imaging parameters for gSlider-SMS acquisition protocol. (b) Diffusion directions of the datasets. The 420 directions at b = 1000 s/mm^2^ is a subset of the 840 directions for b = 2500 s/mm^2^ (shown in blue).

To achieve good SNR and high angular resolution, a total of 2808 volumes of dMRI data were acquired across the 9 sessions, consisting of 144 b = 0 s/mm^2^ images, 420 b = 1000 s/mm^2^ diffusion-weighted images (DWIs) and 840 b = 2500 s/mm^2^ DWIs, along with their paired reversed phase-encoding volumes. The two shells at b = 1000 s/mm^2^ and 2500 s/mm^2^ are shown in Figure 2b, where the 420 directions at b = 1000 s/mm^2^ represented a subset of the 840 directions at b = 2500 s/mm^2^. The diffusion vectors were designed using the algorithm described in Emmanuel Caruyer *et al.*^48^ (publicly available tool at http://www.emmanuelcaruyer.com/q-space-sampling.php), which not only provides uniform angular coverage for each shell and for overall multishell sampling, but is also able to provide uniform incremental angular distribution. The TEs of b = 1000 s/mm^2^ and b = 2500 s/mm^2^ were matched. In each session, 312 dMRI volumes were acquired, including 280 DWIs (140 unique diffusion directions with reversed phase-encoding acquisition) and 32 b = 0 s/mm^2^ images (interleaved with every 10 diffusion images). These data were further divided and acquired in four acquisition blocks, so the calibration data could be updated every ~25 mins and the paired reversed phase-encoding volumes could be interleaved.

In each session, a low-resolution field map and a structural scan (T_1_ or T_2_-weighted image) were also acquired as time permitted. The low-resolution field maps were acquired using a two-TE gradient echo field mapping sequence: TR = 351 ms; TEs = 4.14, 6.6 ms; FOV = 220 × 220 mm^2^; voxel size = 2 × 2 mm^2^; number of slices = 38; slice thickness = 2.5 mm; slice gap = 1.3 mm; flip angle = 45°; bandwidth = 500 Hz/Pixel; total acquisition time = 79 s. The T_1_-weighted images were acquired with the Multi-echo Magnetization-Prepared Rapid Acquisition Gradient Echo (MEMPRAGE)^49^ sequence at both 1-mm isotropic and 0.7-mm isotropic resolutions. The 1-mm MEMPRAGE protocol was acquired using the following parameters: TR = 2530 ms; TI = 1100 ms; number of echoes = 4; TEs = 1.61, 3.47, 5.34, 7.19 ms; flip angle = 7°; FOV = 256 × 256 × 208 mm^3^; bandwidth = 650 Hz/Pixel; GRAPPA factor along phase encoding (PE) direction = 2; total acquisition time = 6 minutes. The 0.7-mm MEMPRAGE protocol: TR = 2530 ms; TI = 1110 ms; number of echoes = 4; TE = 1.29, 3.08, 4.83, 6.58 ms; flip angle = 7°; FOV = 256 × 256 × 146 mm^3^; bandwidth = 760 Hz/Pixel; GRAPPA factor along PE = 2; total acquisition time = 8.38 minutes. T_2_-weighted data were acquired with a T_2_-SPACE^50^ sequence at 0.7-mm isotropic resolution with the following parameters: TR/TE = 3200/563 ms; FOV = 224 × 224 × 180 mm^3^; bandwidth = 745 Hz/Pixel; GRAPPA factor along PE = 2; total acquisition time = 8.4 minutes. All T_1_- and T_2_-weighted data were acquired with image readout in the head-foot (HF) direction, phase-encoding (PE) in the anterior-posterior (AP) direction, and partition in the left-right (LR) direction. One 1-mm isotropic resolution MEMPRAGE, three 0.7-mm isotropic resolution MEMPRAGE and three 0.7-mm isotropic resolution T_2_-SPACE images were acquired in session 3-9, respectively.

### Data reconstruction

The image reconstruction for dMRI was performed on raw k-space data in MATLAB (Mathworks, Natick, MA, USA) (Fig. 1c). First, dual-polarity-GRAPPA^44,45^ (DPG) was used for parallel imaging reconstruction of both in-plane and multi-band accelerations. DPG has been validated to reduce Nyquist ghost artifacts of EPI effectively by correcting for higher-order phase errors^44,45^. Subsequently, POCS^51^ was used for partial Fourier reconstruction to recover high spatial information from partially acquired k-space data. After DPG and POCS, diffusion background phase-corruptions were removed using the real-valued diffusion algorithm to provide real-value data and avoid magnitude bias in subsequent post-processing steps^52^. Finally, gSlider’s slab-to-slice reconstruction^40,41^ was performed to reconstruct thick RF-encoded slabs (3.8 mm) into thin slices (0.76 mm), where T1 recovery was incorporated into the reconstruction model to mitigate slab-boundary artifacts^53^.

The image reconstruction of the T_1_- and T_2_-weighted images and field maps was performed by the vendor’s (Siemens Healthcare, Erlangen, Germany) online reconstruction (including vendor’s implementation of parallel imaging reconstruction, fast Fourier transformation, coil combination, root mean square combination of different echo images and calculation of field maps) and was finally stored in Digital Imaging and Communications in Medicine (DICOM) format.

### Data preprocessing

Preprocessing corrections across the reconstructed image series were performed to correct for signal drift, susceptibility, eddy currents, and gradient nonlinearity induced distortions, and subject motion. An optimized pipeline was used where the warp fields for susceptibility, eddy-current distortion, gradient nonlinearity and motion were estimated, combined, and applied to each image volume in a one-step resampling process to ensure minimal smoothing effects due to interpolation^54,55^. The FMRIB Software Library (FSL)^56–58^ was used to perform these estimations and corrections. The preprocessing pipeline is shown in Fig. 1d and described in detail below:

i. First, intensity normalization and signal drift^59,60^ correction were performed. The non-DW images (b = 0 s/mm^2^) were used to normalize the signal intensity of the data acquired within and across the different acquisition blocks (~25 minutes each). The signal drift within each 25-minute acquisition block was corrected using scaling factors obtained from fitting a second-order polynomial on the mean signals of the non-DW images as a function of the scanned non-DW and DW images. Specifically, there were 8 non-DW images interleaved throughout each 25-minute acquisition block (one non-DW volume every 10 DW volumes, 3.2 minutes). The mean signals of the non-DW volumes were used to fit polynomial coefficients using MATLAB’s *polyfit* function, which were then fed into the *polyval* function to obtain scaling factors for DW volumes in between those non-DW volumes. The signal intensity differences across acquisition blocks were normalized to the mean signal intensity of the first session.
ii. Susceptibility-induced and eddy current-induced distortion estimations and corrections were performed using topup^56,61^ and the eddy-current correction toolbox^62–64^ in the FSL^56–58^. The warp fields for each volume were obtained and stored for later use in the combined one-step resampling correction. Note that topup and eddy-current correction were performed for each individual data subset (2 acquisition blocks with paired reversed phase encoding block), such that a field map could be obtain for each data subset to account for field changes due to field drift along time or subject position changes. In addition to updating the field map, the interaction between susceptibility-induced field changes and subject motion within each data block was also considered by using a dynamic correction algorithm^64^ as implemented in the FSL eddy toolbox^62–64^.
iii. The warp field for gradient nonlinearity correction^54,65^ was calculated for the diffusion data. Here, the gradient nonlinearity correction was performed after topup and eddy correction^54^. This is because of the consideration that the voxel displacements due to susceptibility and eddy current distortions are much larger than those due to within-block rigid motion. Therefore, to obtain better estimation of these larger voxel displacements, the original phase-encoding direction was maintained prior to the estimation and correction for susceptibility and eddy current-induced distortions^54^. The effects of performing gradient-nonlinearity at a later step on the accuracy of within-block rigid motion estimation were considered to be small, because we assume that the confined motion within each acquisition block would only cause small differences in image deformation caused by gradient non-linearity.
iv. The next step estimated larger motion parameters across different data subsets such that the images from different acquisition blocks and sessions could be registered and aligned. This was performed using FLIRT^66–68^ in FSL^56–58^ with 6 degrees of freedom (DOF) rigid motion.
v. Finally, the warp fields for susceptibility, eddy-current distortion, gradient nonlinearity and motion from steps ii-iv were combined and applied to each image volume in a one-step resampling process to ensure minimal smoothing effects due to interpolation^54,55^. The Jacobian matrix computed from the combined warp field was used to correct for the signal intensity affected by the transformation.

All 18 data subsets from 9 sessions and their corresponding b-value and b-vector files were concatenated together and stored in compressed Neuroimaging Informatics Technology Initiative (NIfTI) format for downloading (Fig. 1d, data output B) and further analysis. Freesurfer^69^ functions were used to load and save NIfTI data.

To provide flexibility for the end users to use their own preprocessing pipeline of choice, minimally preprocessed data are also made available for download in compressed NIfTI format (Fig. 1c, data output A). With the minimally preprocessed data, only the gradient nonlinearity correction has been performed without any other preprocessing corrections. The rationale in performing this correction is that the gradient nonlinearity coefficients needed in this correction are considered proprietary information by Siemens, and are not publicly available.

T_1_ and T_2_-weighted images are available for download in compressed NIfTI format (Fig. 1c, data output A). After online reconstruction, the DICOM data were converted to NIfTI format using dcm2nii^70^ tool of MRIcron (http://www.mricro.com), and MRI gradient nonlinearity correction was performed. Subsequently, face and ear regions were masked to de-identify the high-resolution structural images using the defacing tool provided by FSL^56–58^. The averaged images of the three 0.7-mm T_1_ and T_2_-weighted images are also provided after registration, averaging and de-identification. The low-resolution field maps are provided after gradient nonlinearity correction.

### Data Records

All data records listed in this section are publicly available in dryad^71,72^.

#### dMRI data

i. The preprocessed dMRI data are saved under ‘preprocessedDWIData.zip’^71^. All the images in 9 sessions are saved in a single NIfTI file. The corresponding b-value and b-vector files are also provided.
ii. The minimally preprocessed dMRI data are saved under ‘reconDWIData.zip’^72^. 18 individual data subsets are saved in the compressed NIfTI format (paired phase encoding direction concatenated together), encoded by labels identifying scan session and data subset number. The corresponding b-value and b-vector files are also provided in the folder.

#### Structural data

i. T_1_-weighted images are saved under ‘T1.zip’^71^ in compressed NIfTI format, containing one 1-mm MEMPRAGE image (echo combined), three 0.7-mm MEMPRAGE images (echo combined) and their average image.
ii. T_2_-weighted images are saved under ‘T2.zip’^71^ in compressed NIfTI format, containing three 0.7-mm T2-SPACE images and their average image.
iii. Field maps for all 9 sessions are saved under ‘FieldMap.zip’^71^ in compressed NIfTI format, labeled by numbers identifying scan session.

Supplementary to the above data, a readme text file^71^ is included in the repository with more information of the datasets.

### Technical Validation

#### Assessment of DW image quality and head motion

The single and mean DWIs at b = 1000 s/mm^2^ and b = 2500 s/mm^2^ in three orthogonal views are shown in Figure 3a, which highlight the high image quality of this isotropic resolution dataset in all three spatial orientations with minimal reconstruction and ghosting artifacts. The high isotropic spatial resolution of the data reveals detailed structures of the brain (e.g., thalamus, globus pallidus, cerebellum) and sharp delineation of the gray-white matter boundaries (apparent in mean DWIs).

**Figure 3.**
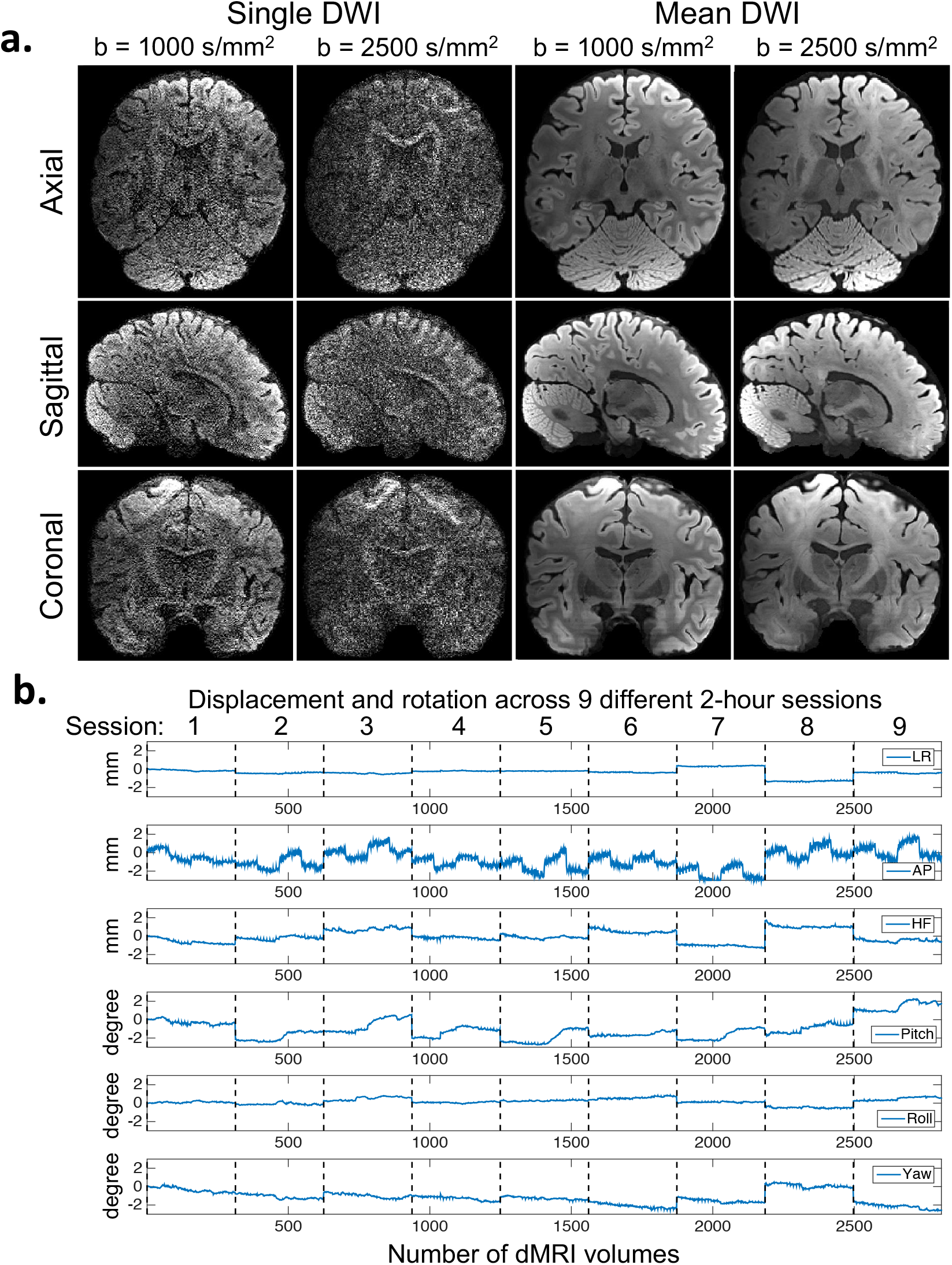
DW images and subject movement measurements. (a) Single DWI and mean DWI images at different b-values shown in three orthogonal views. (b) subject movement measurements of all 2808 dMRI volumes acquired in 9 two-hour sessions. From top to bottom are translations along left-right (LR), anterior-posterior (AP), head-foot (HF) and rotations around LR (Pitch), AP (Roll), HF (Yaw) axis. The motion parameters were estimated relative to the first b = 0 s/mm^2^ volume in the first session. dMRI volumes acquired in different sessions on different dates are separated by black dashed lines. As shown, both within-session movements and across-session head position changes were kept small with the use of the personalized motion stabilizer.

Subject motion between the 2808 dMRI volumes across the 9 two-hour sessions was evaluated in Figure 3b. The motion parameter estimates from different sessions acquired on different dates were concatenated together in a chronological order and separated by dashed lines. Within each two-hour session, the translations along the LR and the HF directions as well as the rotations around the AP (roll) and the HF axes (yaw) were all confined within a ±1 mm/degree range. The rotations around the LR axis was slight larger but was still within a ±2-degree range. The estimates of translational motion along the AP direction appeared to be the highest and contained fluctuations and large jumps between the 25-minute acquisition blocks with opposing phase-encoding directions. This likely resulted from the inclusion of some eddy-current induced translation along the phase-encoding direction, which is known to be difficult to differentiate from actual subject motion since they affect the data in the same way. However, whether this translation is included in the eddy current-induced estimate or the subject movement-induced estimate will not affect the image motion correction. Across different scan sessions, the abrupt changes in motion parameters represented differences in head position placements across sessions. With the use of the personalized head motion stabilizer, the position changes across different sessions were minimized to within ±2 mm/degrees.

#### Assessment of diffusion modelling outcomes of the multi-session dataset

The quality of the multi-session dataset was also assessed by evaluating the diffusion modelling outcomes. Specifically, the diffusion tensor (DT) model and a multi-shell extension of the ball & stick model^73–75^ implemented in the bedpostx tool in FSL were used. The necessity and benefits of the multi-session data acquisition were illustrated by comparing the above diffusion outcomes obtained using data from 1, 3 and 9 sessions (total dMRI data acquisition time of 1.6, 4.8 and 14.5 hours). As shown in Figure 4, increasing the amount of data significantly increased the SNR and quality of the fractional anisotropy (FA) maps. The FA maps obtained using data from all 9 sessions revealed exquisite detail within fine-scale structures, including the radiating fiber bundles within the corpus callosum, highlighted as red stripes on the FA map (Fig. 4, zoomed-in area, arrow-A); the ~1-voxel-width line of reduced-FA separating the corticospinal tract from an adjacent tract (Fig. 4, zoomed-in area, arrow-B), highlighting the low level of blurring and high spatial accuracy of this data acquired across 9 two-hour sessions; and the gray matter bridges that span the internal capsule, giving rise to the characteristic stripes seen in the striatum (Fig. 4, zoomed-in area, arrow-C). In addition, the increased SNR and the denser q-space sampling from the multi-session data dramatically increased the ability to resolve three-way crossing fibers (see Fig. 4 for increased volume fraction of the third fiber shown in red-yellow on top left) and revealed complex intermingling fiber bundles within the centrum semiovale (see Fig. 4 fiber orientation maps modelling 3 fiber compartments per voxel), where the callosal fibers, the corticospinal tract and the superior longitudinal fasciculus intersect.

**Figure 4.**
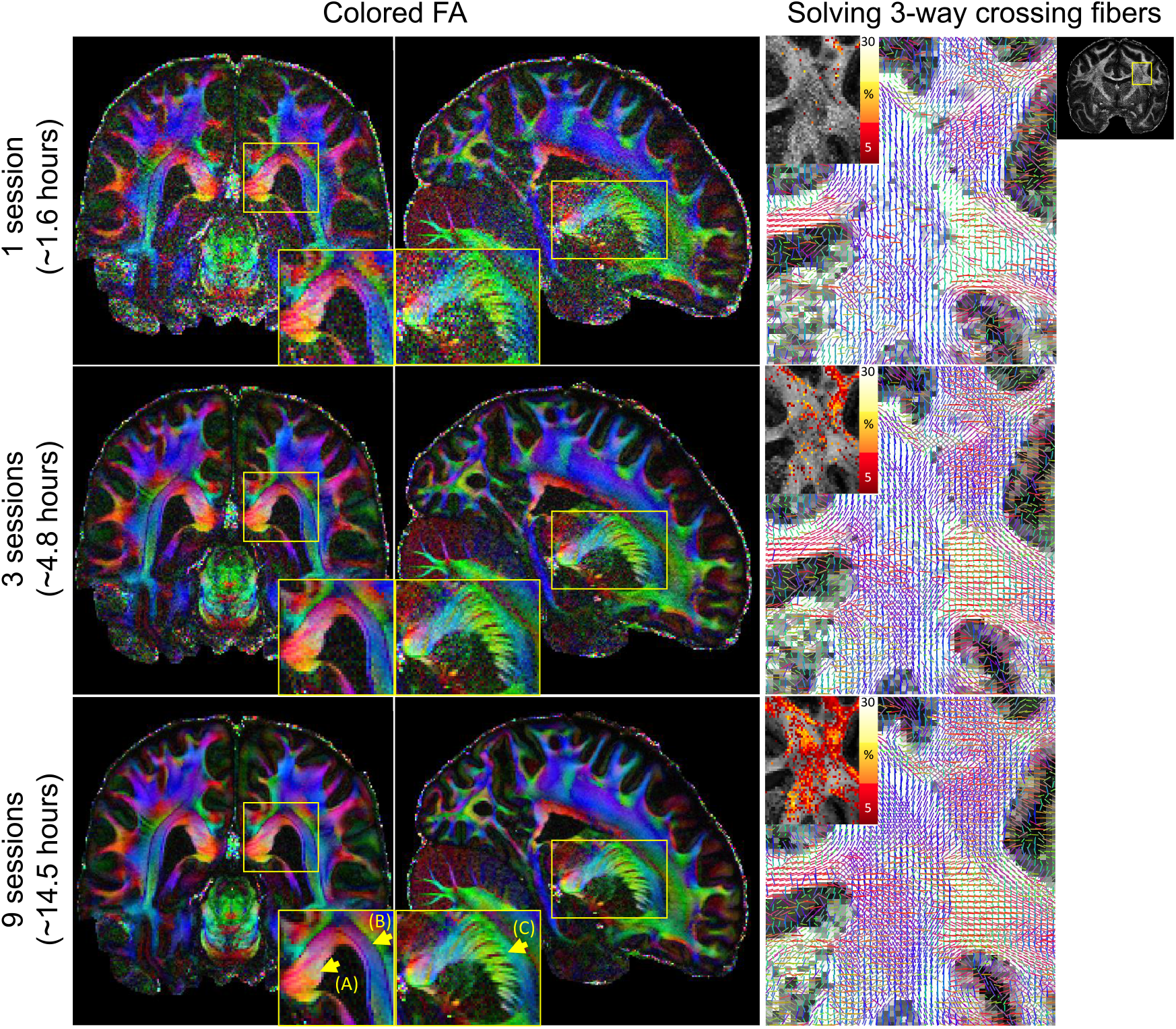
Comparison of FA maps and crossing fiber analysis results using data from 1, 3 and 9 acquisition sessions. On the left, colored FA maps in three orthogonal views are shown for data acquired in 1 session (top), 3 sessions (middle) and 9 sessions (bottom). The 9-session data provides high SNR FA and reveals exquisite fine-scale structures as shown in the zoomed-in areas (yellow boxes), including (A) the radiating fiber bundles within the corpus callosum; (B) the 1-voxel-width line of reduced-FA separating the corticospinal tract from an adjacent tract, highlighting the low level of blurring and high spatial accuracy of this data; and (C) the gray matter bridges that span the internal capsule. On the right, zoomed-in fiber orientations (three fiber compartments per voxel) within centrum semiovale (yellow box) overlaid on gray-scale maps of the total anisotropic volume fraction are shown for those data as well. In addition, the volume fraction of the third fiber are shown in red-yellow on the top left of each fiber orientation map to visualize the sensitivity of solving 3-way crossing fibers. Both fiber orientation and third-fiber volume fraction are shown when the respective volume fraction is higher than 5%. The denser q-space sampling and higher SNR of the 9-session data increases its ability to solve three-way crossing fibers and to reveal complex intermingling fiber bundles within the centrum semiovale.

#### Assessment of spatial resolution

A comparison of diffusion results at different spatial resolutions was performed to validate the benefit of using high spatial resolution acquisition. The dMRI data at 0.76 mm isotropic resolution was compared with downsampled data at 1-mm, 1.25-mm and 1.5-mm isotropic resolution (using sinc interpolation on dMRI data, i.e. cropping in the kspace of the dMRI data). Figure 5 shows the colored-FA maps of four zoomed-in areas (yellow squares) at those spatial resolutions. Improved delineation of fiber bundles and more detailed structures can be captured under higher spatial resolution. Figure 6 shows the comparison of DTI tensors at 1.5 mm and 0.76 mm isotropic resolution, which consistently highlights the improved capability of the submillimeter dMRI data to visualize subcortical white matter as it turns into the cortex. A clear dark band with reduced FA at the gray-white junction can be seen at 0.76-mm resolution (red arrows, top and middle rows). In addition, 0.76-mm data detected more sharp-turning fibers such as those connecting cortical regions between adjacent gyri (red arrows, bottom row).

**Figure 5.**
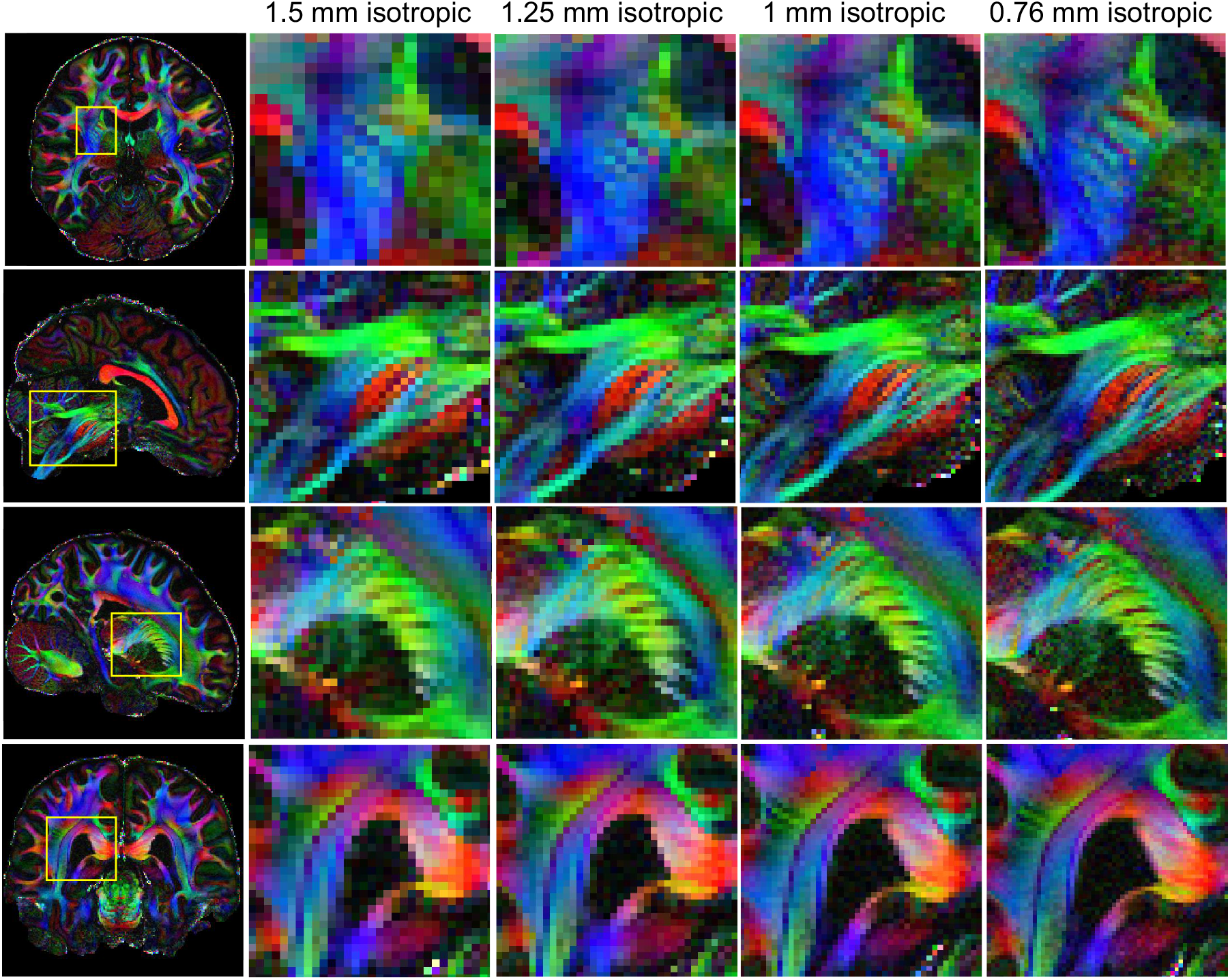
Comparison of dMRI results at different spatial resolutions. (a) Colored FA maps and their zoomed-ins at different brain regions (yellow boxes) and at different isotropic resolutions.

**Figure 6.**
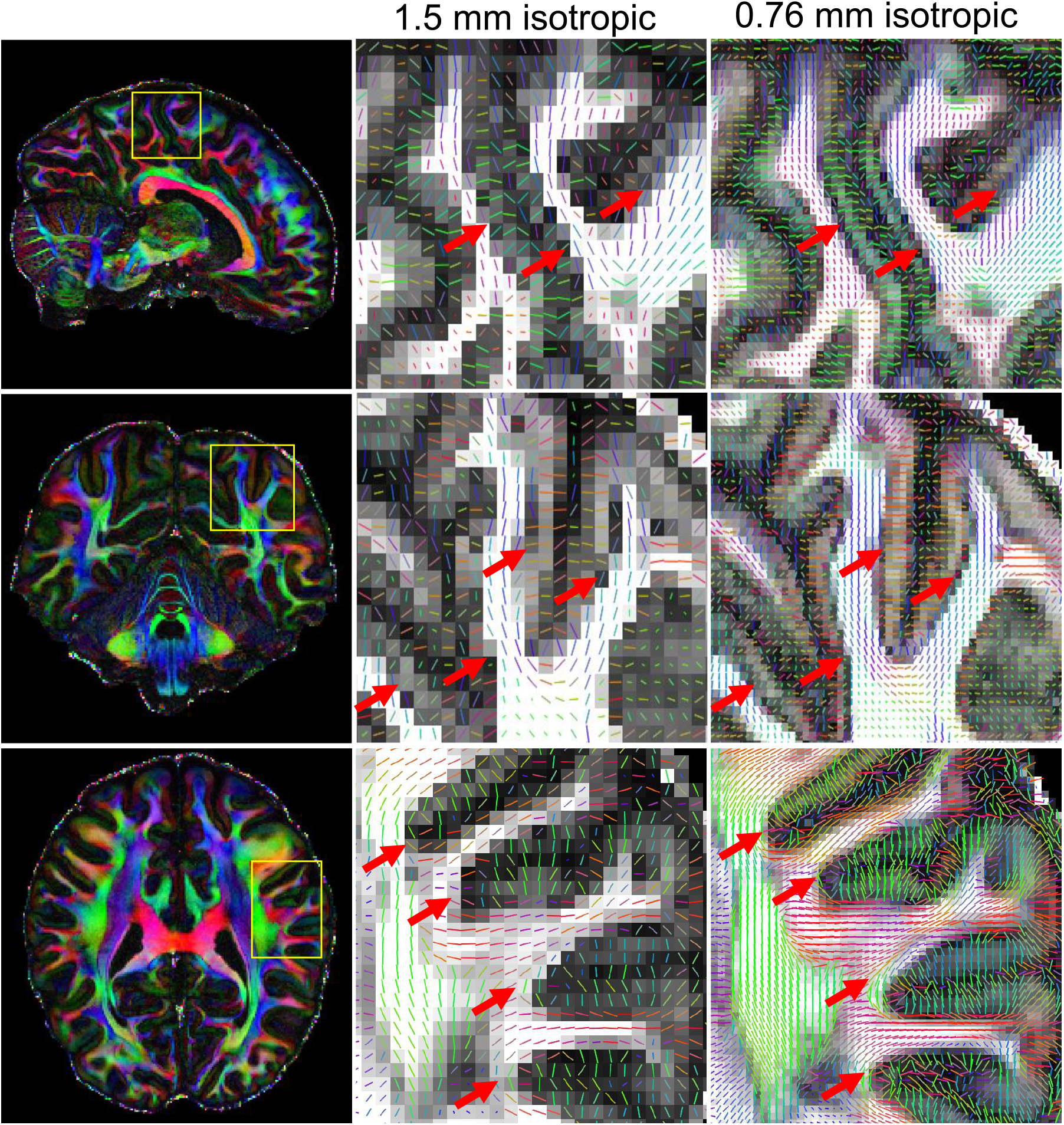
Comparison of DTI principal directions at 1.5-mm and 0.76-mm resolution. Zoomed-in FA and DTI principal vector maps are shown in the area highlighted by yellow boxes. High resolution dMRI data shows improved ability to visualize subcortical white matters as it turns into the cortex. A clear dark-band with reduced FA at the grey-white junction can be seen (red arrows, top and middle rows) at 0.76 mm resolution, and more sharping-turning fibers such as those connecting cortical regions between adjacent gyri (red arrows, bottom row) can be detected.

## Code Availability

The MATLAB code for the preprocessing steps described above are included in the released dataset. The code make use of the functions in the FSL^56–58^ toolbox and Freesurfer^69^ that are freely available for download at https://fsl.fmrib.ox.ac.uk/fsl/fslwiki/FslInstallation and https://surfer.nmr.mgh.harvard.edu/fswiki/DownloadAndInstall.

## Acknowledgements

This work was supported by the NIH (R01-EB020613, R01-EB019437, R01-MH116173, U01-EB025162, and U01-EB026996, U01-MH093765, P41-EB015896, P41-EB030006, K23-NS096056) and NIH shared instrumentation grants (S10-RR023401, S10-RR023043, and S10-RR019307). We thank Dr. Cornelius Eichner for sharing his python-based code for optimized preprocessing of dMRI data with one-step resampling.

## Author contributions

F.W. conceptualized and conducted the study, acquired and processed the data, and wrote the manuscript. Z.D. helped with the data acquisition and the data processing, and contributed to the manuscript. Q.T., C.L., and Q.F. helped with the data processing. W.S.H. helped with the DPG reconstruction. B.K. provided the 64-channel phased-array coil. J.R.P., L.L.W., and S.Y.H. provided the conceptual discussion and contributed to the manuscript. K.S. conceptualized and supervised the study, and contributed to the manuscript.

